# Octopamine and tyramine dynamics predict learning rate phenotypes during associative conditioning in honey bees

**DOI:** 10.1101/2025.08.25.672077

**Authors:** L. Paul Sands, Hong Lei, Seth R. Batten, Alec Hartle, Terry Lohrenz, Leonardo Barbosa, Dan Bang, Peter Dayan, Matt Howe, Brian H. Smith, P. Read Montague

**Affiliations:** Fralin Biomedical Research Institute, Virginia Tech; School of Life Sciences, Arizona State University; Center of Functionally Integrative Neuroscience, Aarhus University, Aarhus, Denmark; Max Planck Institute for Biological Cybernetics, Tubingen, Germany; University of Tubingen, Tubingen, Germany; Department of Physics, Virginia Tech

## Abstract

Biogenic amines are fundamental for physiological homeostasis and behavioral control in both vertebrates and invertebrates. Monoamine neurotransmitters released in target brain regions conjointly regulate adaptive learning and plasticity. However, our understanding of these multi-analyte mechanisms remains nascent, in part due to limitations in measurement technology. Here, during associative conditioning in honey bees, we concurrently tracked sub-second fluctuations in octopamine, tyramine, dopamine, and serotonin in the antennal lobe, where plasticity influences odorant representations. By repeatedly pairing an odorant with subsequent sucrose delivery, we observed individual differences in the conditioned response to odor, which occurred after a variable number of pairings (learners) or not at all (nonlearners). The distinction between learners and non-learners was reflected in neurotransmitter responses across experimental conditions. Remarkably, the speed of learning – the number of pairings prior to a proboscis extension reflex – could be predicted from monoamine opponent signaling (octopamine–tyramine), from both the first presentation of the odorant alone, prior to any pairing with sucrose, and from the first conditioned response to the odorant, coming after a number of sucrose pairings. These results suggest monoamine signaling phenotypes may relate directly to the now widely-reported socially-relevant genetic differences in honey bee learning.

## Main Text

Honey bee foragers are remarkably good at learning the associations between floral cues and the nectar or pollen rewards that their colonies need in order to survive in rapidly changing landscapes(*1*). As a result, honey bees have been developed as a model for understanding cognition in natural and laboratory settings(*2*). Nonassociative and associative conditioning in honey bees are typically studied in a controlled setting using the Proboscis Extension Response (PER) conditioning procedure, which involves pairing an odor with sucrose reinforcement such that the bees develop associatively conditioned responses to the odor(*3*). Classical conditioning of the PER can occur quickly, with some bees only taking a few pairings of an odor conditioned stimulus with food reinforcement, and the memory is retained for several days thereafter. However, individual bees differ in whether or how fast they learn(*4*), and individual differences revealed in PER are correlated with worker foraging decisions in a colony-level foraging strategy(*5*).

PER conditioning has been used successfully to uncover neural correlates of the cellular and molecular substrates of memory(*2, 6*). Using this behavioral protocol, different biogenic amines have been implicated in driving excitatory or inhibitory neural plasticity underlying the behavior. For instance, pioneering work in the 1990’s demonstrated a role for the monoamine octopamine in mediating honeybee PER conditioning, showing that electrophysiological excitation of the octopamine-releasing interneuron VUMmx1 of the subesophogeal ganglion could substitute as a reinforcement signal during associative learning(*7, 8*). Still, several different biogenic amines typically affect processing within subnetworks of the brain; for example, the VUMmx1 neuron must make tyramine in order to make octopamine, and it probably releases both neurotransmitters in the antennal lobes and mushroom bodies(*9, 10*). However, to date, it has remained difficult to further resolve the mechanistic roles of multiple biogenic amines during PER conditioning, due in part to a lack of direct, moment-to-moment estimates of neurotransmitter dynamics in the honey bee brain during associative learning.

Here, we used a machine learning-augmented electrochemistry method (voltammetry), which has been implemented in humans(*11–15*) and validated *in vivo* in rodents(*16*), to concurrently extract sub-second estimates of four neurotransmitters important for honey bee sensory processing and learning: dopamine (DA), serotonin (5HT), tyramine (TYM), and octopamine (OA). These monoamines play integral roles in physiological homeostasis, movement, and adaptive learning processes in invertebrates, with OA and TYM commonly understood as the invertebrate analogs to epinephrine and norepinephrine in the mammalian adrenergic system(*17*). We deployed this method in the antenna lobe of the brain during a standard PER odorant conditioning task, where the development of a PER is the behavioral readout of the conditioned response. The antennal lobes are the site of first-order synaptic interactions between olfactory sensory neurons from the antennae in the periphery and interneurons of the brain(*18*). Associative and non-associative behavioral conditioning change the neural representations of odors in the antennal lobe to increase separation of odors that bees need to discriminate (*19–21*), at least in part through action of octopamine(*22*). Moreover, there is now evidence that OA and TYM may correlate, respectively, with excitatory(*8, 22*) and inhibitory(*23*) behavioral conditioning and the foraging strategies they support (*24*). However, more generally, the nature of the phasic signaling activity and the potential computational roles of these various monoamines during honey bee associative learning remain unknown. Here we show that OA and TYM opponent signalling could regulate a state in honey bees that affects how individuals process information.

## Results

We sought to record concurrent, sub-second changes in four monoamines during PER odorant conditioning. Fig. 1A and 1B shows the *in vitro* calibration protocol and performance of the electrochemistry method, which uses an ensemble of cross-validated deep convolutional neural network (CNN) models to generate monoamine concentration estimates from *in vivo* electrochemical recordings during honeybee behavioral conditioning. Fig. 1C and 1D show the elements of the conditioning paradigm and the behavioral measures that categorized bees as learners or non-learners based on development of a PER. A total of 18 bees were involved in our analyzed sample (see Methods for more details on bee selection); of those 18 bees, 10 developed a PER (‘learners’) after repeated pairings of an initially novel odorant (hexanol) with sucrose, and 8 responded to the sucrose US but never showed a PER to odor within the timeframe of the experiment (‘nonlearners’). The PER behavior itself can be treated as an all-or-none response in which the proboscis is extended in response to an odor or to antennal stimulation with a sucrose solution (see Methods). With repeated pairings of sucrose with an odor conditioned stimulus, the PER will occur in response to the odor alone(*3*). Within the learners, an individual bee’s learning rate, defined as the number of odor-sucrose pairings before the first extension of the proboscis to the conditioned odorant prior to delivery of sucrose reinforcement, varied between 3 and 8 pairings (Fig. 1E).

**Fig. 1.**
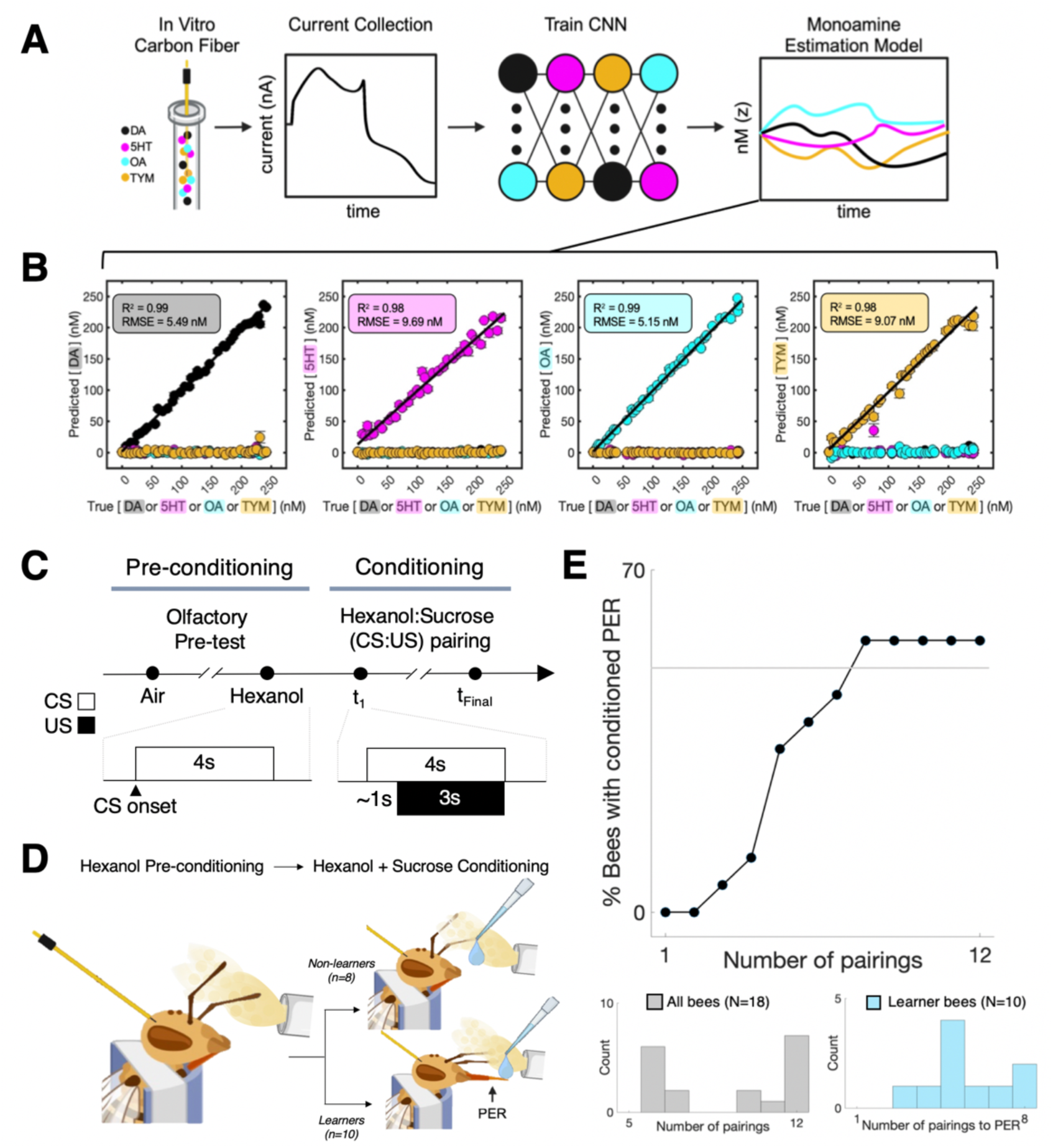
Monoamine vector estimation during honeybee odorant conditioning. (A) Diagram of machine-learning enhanced voltammetry procedure for monoamine concentration prediction. We constructed carbon fiber electrodes and exposed them to varying concentrations of dopamine (DA), serotonin (5HT), octopamine (OA), and tyramine (TYM) against a background of varying pH while using fast scan cyclic voltammetry to measure current responses to each combination. These responses were then used to train an ensemble of deep convolutional neural networks (CNN) via cross-validation to generate a multi-analyte concentration estimation model. (B) Performance of the trained ensemble. Results shown for out-of-sample predictions (nM) for DA (black), 5HT (magenta), OA (cyan), and TYM (yellow), with each molecule shown with their mean ± SEM defined across the 5 different electrodes using in the experiments. (C) Experimental timeline. Each experiment began with a pre-conditioning phase, consisting of olfactory stimulation trials (air (control) and hexanol), followed by an odorant conditioning phase, consisting of up to twelve hexanol:sucrose (CS:US) pairings. Insets depict the approximate timings of CS and US delivery across trial conditions; note that sucrose (US) delivery during conditioning trials occurred ∼1s after the hexanol puff. (D) After completing the conditioning phase, individual bees were classified as “learner” bees (n=10) if they demonstrated a PER to the CS during conditioning, with the remainder classified as “non-learner” bees (n=8). (E) Population curve of PER learning behaviors across conditioning trials for all bees (N=18); bottom: distribution of the number of conditioning trials across all bees (left) and, for learner bees (right), the number of pairings before having PER to the CS (i.e., learning rate). Note that learner bees each had a variable number of trials from the PER trial to the final pairing.

Overall, across both pre-conditioning odorant exposure and odorant exposure during conditioning trials, we show that neurotransmitter fluctuations differed between learners and nonlearners (tables S1-S3), corroborating the observed behavioral differences between the groups. Therefore, we focused on individual differences in neurotransmitter response dynamics within learners for our primary analyses. For learners, the monoamine dynamics during the very first presentation of odor during the pre-conditioning phase (before any reinforcement), and during the conditioning trial where the bee first demonstrated a conditioned PER to the odor presentation, revealed a remarkable relationship to individual bee learning rates.

During the pre-conditioning phase (Fig. 2), all bees were exposed to a control air stream (fig. S1) and an air stream containing hexanol (fig. S2), which was used as the conditioned stimulus later in the experiment, as well as the other primary alcohols heptanol and 2-octanone on separate trials. These three odorants are among several that have been widely used in honey bee PER conditioning, and the similarities in neural activity they elicit in the Antennal Lobe reflect similarities of the molecular structures(*25*). To investigate the computational roles of biogeneic amines in guiding behavioral adaptations during associative learning (*26*) we analyzed the time series of OA, TYM, DA, and 5HT as well as two opponent pairs: DA–5HT(*27*) and OA–TYM. We evaluated the first pair given known roles for dopamine and serotonin opponency in mammalian behavioral control [20,(*28, 29*); we evaluated the latter pair because of their known involvement of octopamine and tyramine in antagonisitic regulation of insect behaviors(*17*). We computed the response level of each neurotransmitter and opponent pair by integrating the time series values within the 4 second olfactory stimulation period for each condition (i.e., area under the curve, AUC; Fig. 2B).

**Fig. 2.**
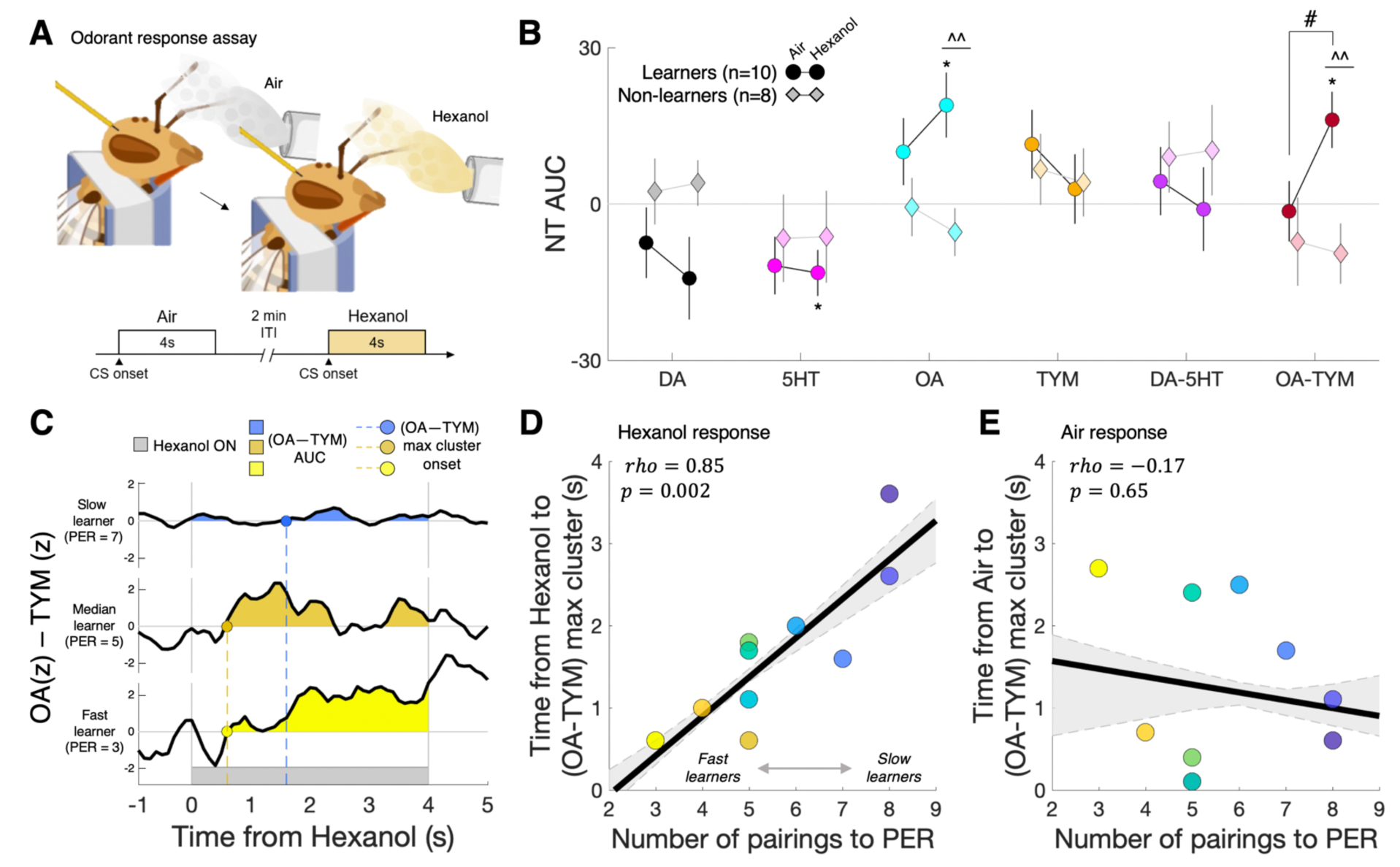
Octopamine and tyramine opponent response to hexanol pre-conditioning predicts PER learning rate. (A) Schematic of honey bee odorant response assay. Bees were exposed to 4 seconds of (control) air and hexanol across two trials, with each trial separated by two minutes. (B) Response levels (AUCs) for neurotransmitters and opponent pairs across the odor stimulation conditions. Circles and diamonds represent mean group values; vertical lines depict SEM. *p<0.05,***p<0.005 one-sample t-test; ^^p<0.01 two-sample t-test; #p<0.05 paired t-test. (C) Example slow, median, and fast learner bees’ (n=10) [OA-TYM] opponent response to hexanol, with positive AUC clusters highlighted and the onset of the maximum positive cluster indicated by the dashed line. (D) Correlation of individual learner bees’ (OA-TYM) max positive cluster onset time during hexanol exposure with their subsequent learning rate (trials to PER) during conditioning. (E) Correlation of learner bees’ (OA-TYM) max positive cluster onset time during control air puff with their learning rate. In both (D,E), the black line indicates the group-level correlation (rho) and the grey dashed lines and shaded region indicate the maximum and minimum of the (k-fold) bootstrapped group-level correlation.

We first tested whether the response levels of biogenic amines or their opponent pairs were related to well-established differences in whether bees show a conditioned reponse to odor after its pairing with reinforcement. Linear modeling assessing the effects of group (learner or nonlearner), odor (air or hexanol), and their interaction on neurotransmitter response levels revealed significant main group effects for DA, OA and (OA–TYM) responses (table S1). For nonlearners, neurotransmitter and opponent pair response levels to air and hexanol were not significantly different from zero, and there were no differences between air and hexanol responses (Fig. 2B; table S2). In contrast, learners demonstrated significant OA, 5HT, and OA–TYM responses to hexanol, with a specific role for the OA–TYM opponency signal, which not only distinguished hexanol responses between learners and nonlearners (table S1) but also distinguished air and hexanol responses within learners (Fig. 2B; table S2). Given this finding, we sought to determine whether the hexanol neurotransmitter response levels were specific to this odor or generalized to other odors. To this end, we examined the representational similarity of air, hexanol, heptanol and 2-octanone neurotransmitter response patterns (i.e., the 6-vector of AUC values; fig. S3). Across all bees, we found significant similarity in monoamine profiles across all three odors (fig. S3), which indicates that changes in neurotransmitter responses during PER conditioning described below are state- or experience-dependent changes rather than being specific to one or another odorant.

We next evaluated whether neurotransmitter response levels to odor prior to conditioning might relate to the degree to which bees learned at different rates during conditioning. To this end, we regressed the learners’ neurotransmitter response levels to hexanol pre-conditioning against bees’ subsequent learning rates and found that the timing and magnitude of the OA–TYM opponent response to odor was predictive of the number of pairings eventually required for the bee to develop a PER (Fig. 2C and 2D; fig. S4). That is, learners that were fast to acquire a PER during conditioning demonstrated both a larger OA–TYM AUC response (fig. S4B) and an earlier onset of the maximum OA–TYM cluster (Fig. 2C and 2D; fig. S4A) in response to odor exposure pre-conditioning as compared to slower learners. These two measures, the OA–TYM response magnitude and timing, were significantly related to each other (fig. S4C), suggesting a step-change-like functional response that was reflected in the group-averaged OA–TYM time series (fig. S2F). Importantly, these measures were specific to odor sensation and not air stimulation (Fig. 2E; fig. S4D). Moreover, we found that this effect was absent for DA-5HT opponency (fig. S5), further emphasizing the specificity of the odor-evoked OA–TYM opponency as predictive of learning rate. Lastly, to investigate how learning rate might relate to latent patterns in the temporal dynamics of the neurotransmitter time series in response to heanol, we performed a singular value decomposition (SVD) of bee monoamine dynamics in response to odor exposure pre-conditioning (fig. S6). This SVD revealed one singular vector with weights reflecting a similar step-change or sign-flip-like response pattern in the OA–TYM opponency signal (but not dopamine/serotonin opponency), the loading of which was predictive of bee learning rate (fig. S6). It is important to emphasize that the encoding of the learning rate carried by the moment-by-moment opponent OA–TYM signal occurred on the first presentation of hexanol and *before* any pairings with sucrose had occurred, which suggests that bees may be in different states before learning that affect whether or how they attend to and learn about stimulus associations.

In light of our findings that neurotransmitter response patterns to odors pre-conditioning were predictive of subsequent PER learning rate, we sought to quantify the adaptations in neurotransmitter responses in learners and nonlearners across conditioning. We investigated differences in monoamine dynamics in learners and non-learners by specifically focusing on neurotransmitter changes across the first six odor:sucrose pairings (all bees had at least 6 pairings) as well as across conditioning stages, consisting of the first odor:sucrose pairing, the trial of the first conditioned PER response (after 3-8 pairings), and the last odor:sucrose pairing (typically 1-3 pairings after PER trial; Fig. 3, fig. S7-S9). Linear modeling revealed significant main effects of group on response levels for DA, 5HT, OA, and TYM (Fig. 3B; table S3). For the learners, progressing from the first pairing to the PER trial and to the last pairing, we observed DA and 5HT decreased across conditioning stages and OA and TYM demonstrated nonmonotonic adaptations across stages. Similar to our findings from the pre-conditioning experiments, linear modeling indicated that the differences from first to last pairing between learners and nonlearners was specific to OA and the OA–TYM responses (table S3), particularly at the start of learning prior to odor-sucrose pairing. Indeed, the representational similarity of neurotransmitter responses across the first six odor:sucrose pairings (fig. S10) revealed that a given learners’ neurotransmitter response pattern at the first pairing was maintained across the initial pairings until the PER had been acquired, around pairings 4-5 on average, after which the neurotransmitter response patterns were no longer significantly correlated with their initial pattern. In comparison, no such changes in neurotransmitter response vectors were observed for non-learners. Given that DA and 5HT reponse levels in learners decreased steadily across conditioning stages (Fig. 3B), the shift in neurotransmitter representational similarity following PER learning appears related to OA and TYM signaling, with these neurotransmitters demonstrating significant changes in reponse levels before versus after the PER conditioned response (Fig. 3B).

**Fig. 3.**
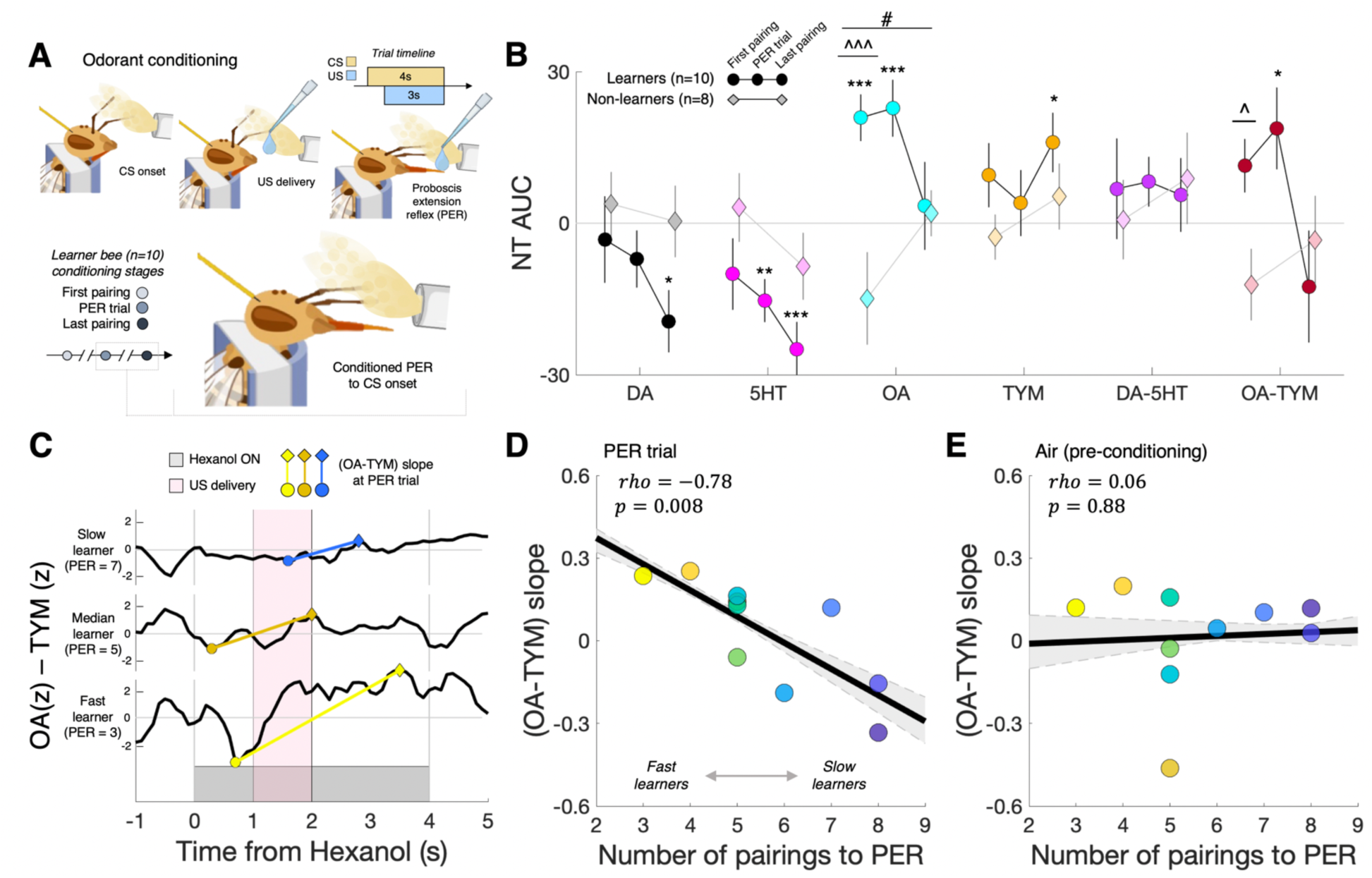
Octopamine and tyramine dynamics during odorant conditioning reflect PER learning rate. (A) Schematic of honey bee odorant conditioning paradigm. On each trial, bees were exposed to 4 seconds of hexanol, with 1.5M sucrose delivered approximately 1 second after hexanol. Each trial was separated by two minutes inter-trial interval. (B) Response levels (AUCs) for neurotransmitters and opponent pairs for the first and last CS:US pairings and, for learners, the CS:US pairing at which a PER was first shown (PER trial). Circles and diamonds represent mean group values; vertical lines depict SEM. *p<0.05, **p<0.01, ***p<0.005 one-sample t-test; ^p<0.05, ^^^p<0.005 two-sample t-test; #p<0.05 paired t-test. (C) Example slow, median, and fast learner bees (n=10) [OA-TYM] opponent signal slope at their PER trial. (D) Correlation of individual learner bees’ (OA-TYM) PER trial slope and their learning rate (trials to PER). (E) Correlation of learner bees’ (OA-TYM) slope during control air puff (pre-conditioning) with their learning rate. In both (D,E), the black line indicates the group-level correlation (rho) and the grey dashed lines and shaded region indicate the maximum and minimum of the (k-fold) bootstrapped group-level correlation.

To investigate the relation of learner bee monoamine response patterns across conditioning stages and individual bee PER learning rate, we quantified the maximum and minimum neurotransmitter responses within the first and second half of each odor:sucrose pairing trial (i.e., roughly before and after sucrose delivery) to give a measure of the rate of change (slope) of each neurotransmitter and opponent pair response across conditioning stages (fig. S11; see Methods). Given the importance of temporal features of learners’ OA–TYM responses to hexanol pre-conditioning in predicting PER learning rate (e.g., step-change onset and size; fig. S4), and the gradually changing similarity in learners’ neurotransmitter response patterns from trial-to-trial (fig. S10), this slope metric is meant to capture individual differences in the magnitude of adaptations in neurotransmitter time series that occur across repeated odor:sucrose pairings. We found that the rate of change of the OA–TYM opponency signal during the first conditioned proboscis extension response (PER trial) was significantly correlated with PER learning rate (Fig. 3C and 3D; fig. S11A), which was specific to odorant conditioning and not observed for pre-conditioning air puff (Fig. 3E). Additionally, the rate of change of the OA–TYM opponent signal at the PER trial was highly correlated with the timing of the OA–TYM response to odor exposure pre-conditioning (fig. S11B), further highlighting a possible role for the temporal dynamics of OA and TYM signaling in establishing olfactory conditioning sensitivity.

## Discussion

Leveraging a machine learning-enhanced approach to concurrent monoamine estimation, we were able to record for the first time in an insect the dynamics of dopamine, serotonin, octopamine, and tyramine during odorant conditioning in honey bees. We segregated honey bees that showed a strong conditioned response to a series of odorant-sucrose pairings and observed – as expected from prior work(*30*) – variability in the number of pairings required for a PER to develop. Overall, the results of the present paper are consistent with the roles that OA and TYM play as the invertebrate analogs to the mammalian use of epinephrine and norepinephrine(*31*). TYR is the direct precurcor to OA, with TYM converted to OA in a single enzymatic step. Thus, the same modulatory neurons – e.g. the VUM neuron(*7*) – make, and likely release, both transmitters in ratios that have been shown in the context of fruit fly locomotion at least to be regulated by factors such as food availability (*32*). Both modulators have also been associated with attractive and repulsive behaviors related to phase change in locusts (*33*) and opponent effects on the flight pattern generator in moths (*34*). Our finding that opponent signaling between OA and TYM on the first presentation of the odorant predicted individual bees’ PER learning rate suggests strongly that opponency profiles at least partially determine “signaling types” that relate to whether and how quickly a honey bee forager responds to new information. Future research could relate this trait to individual bees acting to build more stable statistical models of nectar predictors (slower learners) versus bees more tuned to exploiting current models (faster learners). Such differences across foragers within a colony, which is determined in part by heritable differences among workers(*4, 23*), could allow the colony to more adaptively respond to rapid changes in resource distributions(*35*).

The ability to record vectors of neurotransmitter dynamics will now enable a more thorough evaluation of how biogenic amines are involed in different forms of learning and more generally in regulation of behavior. Here, we have used individual differences in learning rates across bees, commonly observed in PER conditioning(*4, 36*), to test hypotheses about how individual and opponent monoamine signalling can be associated with this behavioral distinction. Further experiments are now possible and warranted using different types of treatments to test how signalling is associated with determining learning-related states or signals of reinforcement directly. These treatments include, for example, presentation of odor(*37*) or sucrose(*38*) alone, contrasting backward and forward pairing(*39*), use of an explictiy unpaired procedure(*40*), and treatments that produce prediction errors(*41*), all of which produce different types of nonassociative, excitatory or inhibitory learning.

Our work also parallels recent findings in mammals that opponent signaling is important for regulating plasticity in the brain. A recent experiment in rodents strongly suggests that DA and 5HT opponency acts as a critical and composite learning signal(*27*). Although the opponency system we describe here prepares a bee for learning new information, we cannot at this point confirm which monoamines, or combinations of them, represent reinforcement, which has been reported to be OA in honey bees(*7*) and DA in fruit flies(*42*). We observed that DA and 5HT response levels in learners during conditioning each decreased linearly across conditioning stages resulting in an opponency signal not different from zero. The decreases in DA and 5-HT hint at a role for these monoamines in associative conditioning that will need to be a focus of future research, but that role appears not to be in opponency signaling as for OA and TYR. Prior work implicating DA in aversive conditioning and punishment signaling in honey bees(*43*) suggests that changes we show here could be seen as a general state-value signal that becomes increasingly positive (i.e., reduction in aversion) across repeated odorant:sucrose pairings during PER conditioning. For OA and TYM, the response levels across learners appeared to change most significantly before versus after the conditioned PER behavior, such that the OA–TYM response related to PER learning rate started positive before the onset of conditioning, remained positive up to the PER trial, and then flipped to negative. This trajectory was driven by concave OA and convex TYM changes that, based on prior work, would be hypothesized to cause decreased appetitive and increased inhibitory conditioning after the PER was learned, respectively, thereby implementing a putative de-sensitization signal that limits ongoing associative learning. In this way, each bee’s OA–TYM signaling type could be tested as not just phenotypically setting a bee’s learning rate pre-conditioning but also regulating explore-exploit choice policies during foraging via differential expression of sensitiziation and habitiuation processes. Ultimately, while further experiments are required to address these outstanding questions, our results are important steps towards a more comprehensive analysis of the roles of octopamine, tyramine, dopamine and serotonin in driving attention and reinforcement learning across species.

## Author contributions

B.H.S and P.R.M.. conceived the study and designed the experiments. H.L and S.B. collected the behavioral and voltammetry data. L.P.S., A.H., T.L. and L.B. analyzed the data and performed statistical analyses. D.B. P.D. and M.H. provided feedback on data analyses and on the manuscript. B.H.S., P.R.M. and L.P.S. wrote the manuscript and developed the figures.

## Supplementary Materials

## Materials and Methods

### Animals

Honey bees (*Apis mellifera*) were raised in the Arizona State University apiary. All pollen foragers (female) were captured at the colony entrance and brought into the laboratory. Each bee was then chilled at 4°C and placed in a plastic harness. Before performing the surgery for neurochemical recordings, bees were tested for motivation by stimulation of their antenna with 1.5 M sucros[14]e– if bees extended their proboscis, they were used in the experiments.

### Neurochemical recording electrodes

Monoamine neurotransmitter recordings were taken from the antenna lobe using in-house made carbon fiber electrodes that were adapted from those described in Kishida et al. [44]. Briefly, these electrodes were made by threading a carbon fiber into a small glass capillary (90 µm OD; Molex). The carbon fiber was then pulled and epoxied such that only ∼150 µm (approximate length of dorsal-ventral axis in a bee) was exposed on one end of the capillary and the carbon fiber at the other end of the capillary was trimmed to ∼2 mm. The ∼2 mm end of the small capillary was then placed into a larger capillary (440 µm OD; Molex #1068450450) containing a Pt/Ir wire such that the carbon fiber overlapped the Pt/Ir wire by ∼1 mm. Silver paint was used to marry the carbon fiber and Pt/Ir wire to enhance electrical conductance. The silver paint was drawn into the large capillary via capillary action before placing the small capillary inside. After a 24 hr curing period, the large and small capillary were then epoxied together using marine epoxy. A gold pin was then soldered on to the Pt/Ir wire exposed on the other end of the large capillary. Some of the gold pin and some of the large capillary were then covered with shrink wrap and epoxy was used to seal the joint. A stainless-steel wire with a gold pin soldered on one end was used as the reference electrode.

### Neurochemical recording equipment and procedures

We obtained voltammetry data in bees using the carbon fiber microelectrodes described above. The voltammetry protocol was based on previous work in humans [11-15, 44] and rodents [45, 46]. The voltage forcing function used was a standard triangular voltage shape repeated at 10 Hz (ramp up from −0.6 V to +1.4 V at 400 V/s, and ramp down from +1.4 V to -0.6 V at 400 V/s [10ms] then hold at -0.6 V for 90 ms). The current response was recorded at a sampling rate of 100 kHz. We used a 97 Hz pre-cycle protocol prior to data collection to aid in microelectrode equilibration. This pre-cycle protocol used the previously mentioned triangular voltage ramp repeated with a shorter holding period. The voltammetry recordings were made using Molecular Devices (Molecular Devices, LLC, San Jose, CA, 95134) equipment. Namely, the Axon Instruments MultiClamp 700B Patch Clamp Amplifier, the Electrochemistry Headstage (CV 7B-EC; slightly modified to allow larger current measures), and the Digidata 1550B Data Acquisition System.

### Neurochemical recordings in the antenna lobe

After the bees were cooled and placed in the plastic stage, their antennae were held in place using Eicosane, a low-melting-point (31^oC^-38^oC^) alkane, and a hole was opened in the head capsule to expose the brain. The in-house made carbon fiber electrode was then placed in the antenna lobe using a stereotaxic frame (Kopf Instruments; Tujunga, CA). Fast scan cyclic voltammetry (FSCV) measurements were then taken, using the equipment and protocols described above, while the bees were exposed to a conditioning procedure (described below). A TTL pulse (AFG1062, Tekronix) was used to align neurochemical and behavioral events.

### Proboscis extension response conditioning protocol

Harnessed bees that responded to sucrose (see above) were put in front of an airflow and odor delivery system. A carbon fiber electrode was then placed in the antenna lobe (see above) and neurochemical measurements were taken during the following procedures. Bees were first presented with an air stream containing 10:1 hexanol on a trial-by-trial basis (all odor presentations lasted 4 seconds). These odor-only trials were meant to serve as controls for the conditioning trials (i.e. to aid in isolating neurochemical signatures related to learning). All of the trials, including the conditioning trials, used a 2-minute intertrial interval (ITI). Bees were conditioned using 10:1 hexanol (conditioned stimulus [CS]) and 1.5 M sucrose (unconditioned stimulus [US]). Hexanol was presented for 4 seconds and sucrose was delivered 1 second after odor onset by gently touching the antennae to elicit proboscis extension that followed by feeding. Once a bee extended its proboscis after the onset of hexanol (PER), it was no longer necessary to touch its antennae. After a PER was established, hexanol, octanone, and heptanol, were presented on separate trials without delivering sucrose. The unreinforced trials served as catch trials to further assess learning. Bees were considered ‘non-learners’ if they did not develop a PER after 12 CS-US pairings, although they always responded to the sucrose-water droplet used as reinforcement.

### Definition of a PER

The honey bee proboscis is made up of several different parts that fold together into a straw-like mouthpart that intakes fluid nectar. It is normally folded underneath the head capsule in a retracted state (Fig 1). When sugar stimulates taste receptors on the antennae or proboscis, or when an odor that has been associated with sugar reinforcement is presented, the first response is that the mandibles open and then the proboscis is extended from underneath the head into a feeding position. Using a fixed reference, such as when the unfolding proboscis breaks the imaginary line between the tips of the open mandibles as observed head on, for example, PER can be quantified as a binary extended-or-not variable on each trial. More degrees of freedom in the PER can be described from electromyographic recordings of head muscles that extend and move the proboscis [47] or by offline analysis of video [48]. A PER can be broken down into eight different parameters that comprise four independent degrees of freedom of movement [47]. Expression of these parameters depends on the nature of the stimulus, e.g. sucrose vs odor (e.g. conditioned odor versus a different odor of varying chemical similarity to the conditioned odor), and the feeding state of the bee (some bees respond more vigorously than others, possibly due to differences in feeding states). Although the PER can differ across bees and test stimuli, for a given bee and test stimulus the PER measured by EMG or video remains fairly constant once the proboscis is extended. For that reason, the binary variable that we use is sufficient for most studies when it reveals differences across response categories, as we show here. However, future use of voltammetry could make use of more detailed measures of the PER as well as information expressed in antennal movements in response to conditioned odors.

### Machine learning electrochemical approach

#### *In vivo* neuromodulator estimates

We estimated *in vivo* neuromodulator concentrations from *in vivo* current traces using an ensemble of deep convolutional neural networks that were trained and tested on *in vitro* data. The *in vitro* data consisted of voltammetry current traces obtained from single 11.5 cm carbon fiber electrodes exposed to known concentrations of dopamine, serotonin, octopamine, tyramine, and pH. We modified the InceptionTime time series classification model [49] to perform multi-variate regression. The model was coded in Python using TensorFlow [50] and Keras. We used equally weighted averages of *in vivo* concentration estimates from multiple InceptionTime models[49].

Specifically, the modified InceptionTime network is based on two ResNet [51] blocks. Each block contains 3 convolutional blocks while each convolutional block is composed of 4 convolutional layers in parallel, each with 32 filters and with increasing kernel sizes– 1, 10, 20, and 40. The output of each of these convolutional layers is stacked together and batch normalization and RELU activations are applied. This output passes through a bottleneck convolutional layer with a kernel of size one, 32 filters, and whose output serves as the input for the next convolutional block. To start the implementation of the ResNet architecture, the input passes through another bottleneck convolutional layer with kernel size 1 and 32 filters and then is added to the output of the first ResNet block. After activation, this serves as the input to the next ResNet block. This operation is repeated for the second ResNet block, except that the input is now replaced by the output of the first ResNet block. Finally, after passing through a global average pooling, the output of the second ResNet block passes through a dense layer with four output nodes– this gives the predictions of the five analytes. All activation functions are RELU, except after the last dense layer, which uses a linear activation function.

All models used the mean squared error loss function. Training schedule used the ADAM optimizer [52], with an initial learning rate of 1e-3 that halved after 5 epochs without decrease of validation loss. These single electrode carbon fiber models did not have a minimum learning rate. The batch size was 64 and the loss on the validation set was calculated after each epoch. The model from the epoch with the lowest validation loss was selected as the final model for that run.

Final predictions of *in-vivo* data are generated using a mixture of experts (i.e. where we train an ensemble of models with the same hyperparameters and average their predictions). Differences in weights generated during initialization, variations in the order in which data is fed to the algorithm during training, and the stochastic descent algorithm all introduce variation to the convergence of each model. Additionally, different validation sets were used in each run to avoid overfitting.

#### *In vitro* training data for carbon fiber bee electrodes

We collected model training data using 5 carbon fiber electrodes, and these electrodes were subsequently used to record *in vivo* current measurements in the bee antenna lobe. Five datasets were collected on each electrode, one for each analyte – dopamine, serotonin, ocotpamine, tyramine – and pH. The data acquisition equipment was of the same type (same component type/number) used in the *in vivo* data collection. Most of the four neurotransmitters (dopamine, serotonin, octopamine, and tyramine) datasets were collected with 40 concentrations of the analyte dispersed at equal intervals over the range from 0 to 250 nanomolar (nM), with a pH of around 6.8 (approximate pH of hemolymph), while the other three analytes were kept at 0 nM. The pH dataset for each electrode was collected with 11 pH values in the range 6.8 to 7.8 and with the concentration of the four neurotransmitters set to 0 nM. For each probe, we randomized the order of neurotransmitter data set collection order, and within each neurotransmitter data set, the order of unique concentrations was also randomized.

To collect data each working and reference electrode was inserted vertically into a 1.5 mL Eppendorf containing the neuromodulator concentration of interest for that collection run. We then collected current data at 10 Hz for 65 seconds using the same measurement protocol deployed during *in vivo* collection. We selected the most stable continuous 15 second section from the second half of the 10 Hz 65 second time window for training in order to reduce variation due to electrical noise and equilibration. See Batten et al., 2024, 2025 for (similar) previously published methods.

#### *In-vitro* model training for carbon fiber bee electrodes

Since we do not yet have enough data collected on 11.5 cm carbon fiber electrodes for a generalized model, we create a specific model for each electrode (e.g. on electrode model). To do this we train a mixture of experts, where the final prediction was the average of an ensemble of 20 training runs. A training run involved a dataset containing 150 sweeps from each unique concentration, that was split into a training set containing 90% of the data and a validation set containing the remaining 10%. The data were split by concentration, such that all data points of any given concentration were in the same set. Before training the model, the data were z-scored within analyte, and then shifted by 10 standard deviations per analyte to avoid zero gradients. After training the model, the inverse of the normalization procedure was performed on the predictions. The models were trained for 100 epochs.

#### *In-vitro* model evaluation for carbon fiber bee electrodes

A 10-fold cross-validation test was performed for each of the 11.5 cm bee electrodes. The *in-vitro* data was split into 10 discrete folds without overlapping values. Each fold was held out as a test set and an ensemble of models was created via a training run on the remaining *in vitro* data. Ensembles of 20 models were used. The mean of the predictions from each 20-model ensemble was calculated for its corresponding test set. All test sets were combined; thus, resulting in all *in-vitro* data points being predicted from ensembles of models that had never been trained or validated on those data points (see Fig. 1B).

#### *In vivo* concentration estimates for carbon fiber bee electrodes

We generated *in vivo* concentration estimates using the same carbon fiber models described above. Note that since we created on-electrode models, we matched *in vitro* models to the carbon fiber that collected the *in vivo* data. *In vivo* current traces were differentiated as in the training and then evaluated by the specific models built for an electrode. Average predictions across this ensemble of models (mixture of experts) were used as the final concentration estimates.

### Neurochemical data analysis

#### Learning rate correlation with monoamine dynamics

Each experiment consisted of the pre-conditioning air and odor stimulation trials and the conditioning trials. For each bee, the model-generated monoamine neurotransmitter time series – 20-second “snippets” centered on the TTL-triggered odor presentation – for every trial were first smoothed using a 500 ms lagging sliding window and then z-scored within-trial (i.e., trial mean and standard deviation computed using full 20-second snippet). Note that this smoothing and normalization were conducted for each neurotransmitter individually, resulting in a 4-vector of monoamine dynamics, which were further cut to a temporal window spanning from 1 second pre-odor to 5 seconds post-odor (odor is present from 0-4 seconds). We computed octopamine– tyramine opponency and dopamine–serotonin opponency time series by taking the difference between the respective neurotransmitter time series pairs after all four neurotransmitters had been individually pre-processed.

Neurotransmitter response levels were defined as the sum of the neurotransmitter time series within the 0-4 second odor presentation window (i.e., taking the area under the curve, AUC). We computed a measure of the rate of change of neurotransmitter time series (i.e., slope) by identifying the magnitude and timing of the maximum or minimum (extrema) of each time series in the 0-2 second window and 2-4 second window of each trial of the pre-conditioning and conditioning phase; the slope metric was then calculated as the difference in extrema values divided by the time between them. To determine the peak opponency response and its relation to learning rate, we identified the timing of positive clusters for each opponent time series via its zero-crossings. We computed the Pearson correlation coefficient for the timing of the onset of the largest positive cluster (i.e., maximum opponent response) and the number of hexanol:sucrose pairings before reaching a PER (i.e., learning rate). We also computed the Pearson correlation coefficient between neurotransmitter response levels (AUCs) and learning rate. This analysis was conducted for all neurotransmitters individually as well as both monoamine opponent pairs, octopamine-tyramine and dopamine-serotonin. We computed a measure of the variation in the correlations between neurotransmitter response features and learning rate via a leave-one-out bootstrapping procedure. For this procedure, we iteratively computed the group-level correlations while holding out each bee once (i.e., k-fold), and we took the minimum and maximum of the resulting set of (k=10) group-level correlations as depicting the variation in the group-level correlation across all learner bees.

#### Representational similarity analysis

We conducted representational similarity analyses (RSA) to characterize the covariation in patterns of neurotransmitter response levels (i.e., AUC values) for learners and nonlearners during the pre-conditioning odor stimulation trials (**SI Appendix, fig. S3**) and across conditiong tirals (**SI Appendix, fig. S10**).

For the pre-conditioning RSA, we first calculated, for each bee individually, the response level (AUC, 0-4s odor window) of each neurotransmitter and opponent pair to the air and odor (hexanol, heptanol, 2-octanone) stimulation trials (**SI Appendix, fig. S3A**). This resulted in a 4-by-6 data matrix (odors-by-NTs) of neurotransmitter AUC values for each bee, from which we then constructed a 4-by-4 representational similarity matrix by computing the pairwise Pearson correlation coefficient of the rows of this data matrix. The lower triangle of this representational similarity matrix captures the covariation in neurotransmitter response patterns for the four pre-conditioning odor stimulation conditions. For instance, the first column of the representational similarity matrix depicts the correlation between a bee’s neurotransmitter response pattern to air stimulation and its neurotransmitter response pattern to the three odors. Looking across all bees, we tested for whether the neurotransmitter response patterns were significantly similar across air and odor conditions by performing a one-sample t-test of bees’ (N=18) Pearson correlation coefficients for each of the seven off-diagonal (lower triangle) element of the representational similarity matrix, with a null hypothesis of corellation coefficients being equal to zero indicating no significant pairwise similarity between conditions at the group level. We performed this pre-conditioning RSA analysis using all bees (N=18) and for learner (n=10) and nonlearner (n=8) bees separately.

For the conditioning RSA, we computed a representational similarity matrix for each bee in a similar manner as the pre-conditioning RSA, though involving the first six odor:sucrose conditioning trials. Thus, for each bee, we first computed a 6-by-6 data matrix (trials-by-NTs) of neurotransmitter AUC values and then computed the individual and group-level representational similarity matrices that capture the covariation in neuroatransmitter response patterns across the six initial odor:sucrose pairings. We chose to examine the first six trials since every bee had at least 6 pairings of odor:sucrose. With this setup, the first column of a bee’s representational similarity matrix depicts the correlation between the neurotransmitter response pattern to the first odor:sucrose pairing and the neurotransmitter response patterns of the ensuing pairings (elements of arrow (i) in **SI Appendix, fig. S10B**). Additionally, the first lower diagonal of the representational similarity matrix (i.e., the diagonal directly below the primary diagonal) depicts the covariation in neurotransmitter response patterns from trial-to-trial (elements of arrow (ii) in **SI Appendix, fig. S10B**). In addition to computing the statistical significance of each off-diagonal element via one-sample t-test, we also computed the significance of the average correlation across the elements of the first column and first lower diagonal of the representational similarity matrix using a one-sample t-test. We conducted this conditioning RSA procedure for all bees and for learners and nonlearners separately; differences between learners and non-learners were computed using a two-sample t-test.

#### Singular value decomposition analysis

We performed a data-driven reduction of honeybee monoamine dynamics using the singular value decomposition (**SI Appendix, fig. S6**). The SVD analysis included all learner bees (n=10) and non-learners (n=8). For each bee, the dopamine, serotonin, octopamine, and tyramine time series, aligned to the onset of hexanol, were concatenated length-wise, converting each bee’s 4-vector of monoamine time series (4 x 61 matrix) into a 1-vector (1 x 4*61). The resulting 1-vector time series were concatenated vertically across all bees, forming an 18x244 matrix **Y**. Computing the SVD of this data matrix **Y** produces a matrix of right singular vectors **V** that represent latent monoamine patterns across bees and a matrix of left singular vectors **U** that reflect the “loading” of the latent monoamine patterns for each bee. We computed the Pearson correlation coefficients of the left singular vector loadings with the learning rate, which revealed one latent pattern, right singular vector 4, which demonstrated a significant correlation. We depicted the monoamine signals of this latent pattern, as well as the dopamine/serotonin and octopamine/tyramine opponent dynamics, by reconstructing the 4-vector of monoamine dynamics based on this latent pattern (i.e., dividing right singular vector 4 back into its component signals; SI Appendix, fig. S6C,D).

#### Statistical modeling and quantitative analysis

We performed a set of linear modeling analyses to quantify the effects of group (learner or nonlearner), odor (air or hexanol), conditioning trial (first or last), and interactions among these variables on neurotransmitter response levels (i.e., AUCs). For the pre-conditioning phase, each linear model consisted of regressing the main effects of group, odor, and their interaction on individual neurotransmitter response levels (i.e., a single model for each neurotransmitter; **SI Appendix, Table S1**); we performed this analysis across all bees (N=18; **SI Appendix, Table S1**) and for learners (n=10) and nonlearners (n=8) separately (i.e., only including main effect of odor; **SI Appendix, Table S2**). Linear modeling was conducted using MATLAB’s function *fitlme* with subsequent ANOVA F-tests over fixed-effects model terms using MATLAB’s function *anova*. We repeated this linear modeling analysis for the conditioning phase, with each model consisting of the main effects of group, conditioning trial, and their interaction (SI Appendix, Table S3).

---

**Fig. S1.**
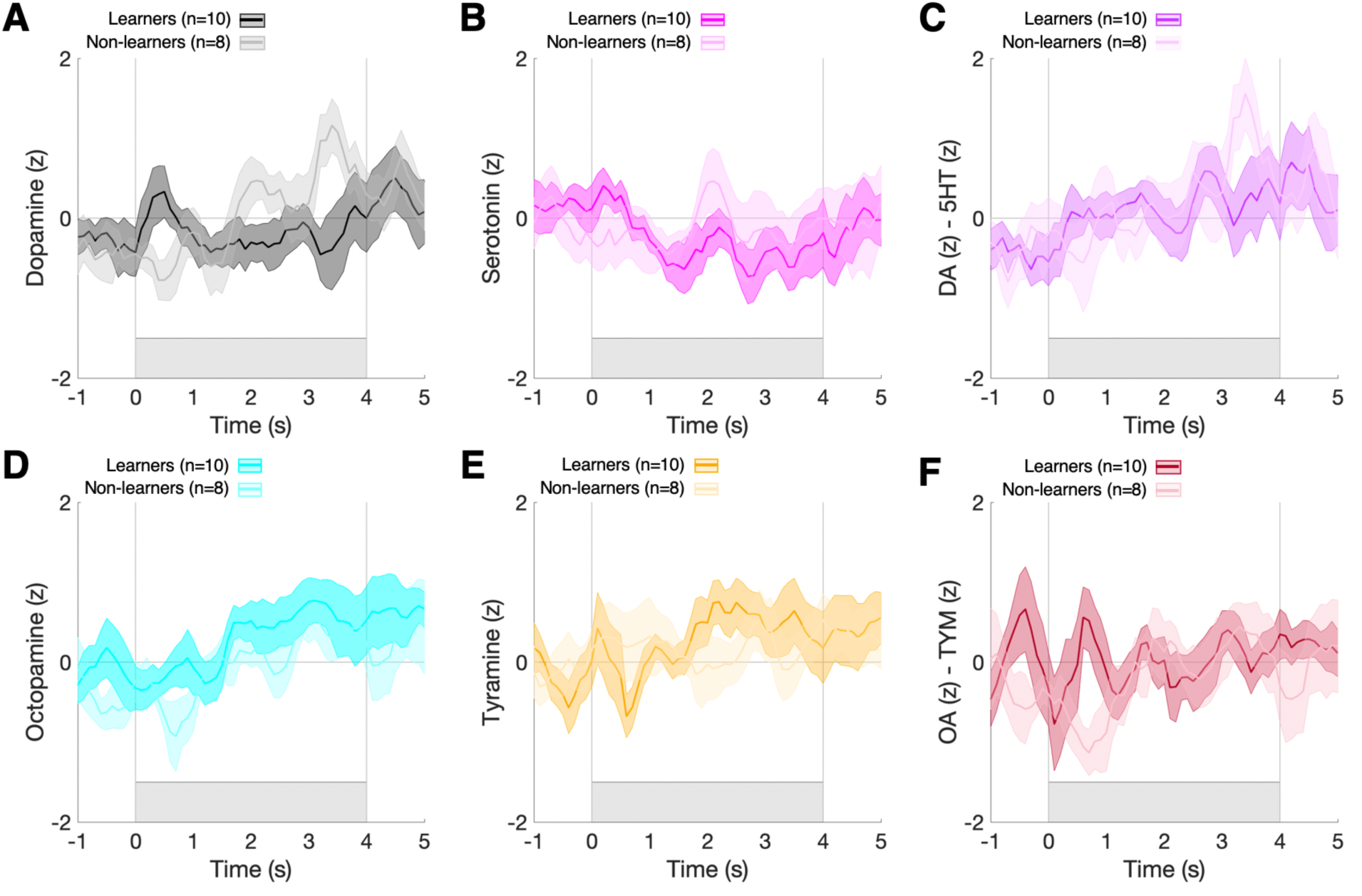
Learner and non-learner bee neurotransmitter response to air puff exposure pre-conditioning. Group-average neurotransmitter and opponent pair time series (10 Hz) in response to air puff pre-conditioning for learner bees (n=10) and non-learner bees (n=8). Shaded regions indicate standard error of the mean (SEM), and the grey vertical lines and shaded rectangle indicate the air puff temporal window.

**Fig. S2.**
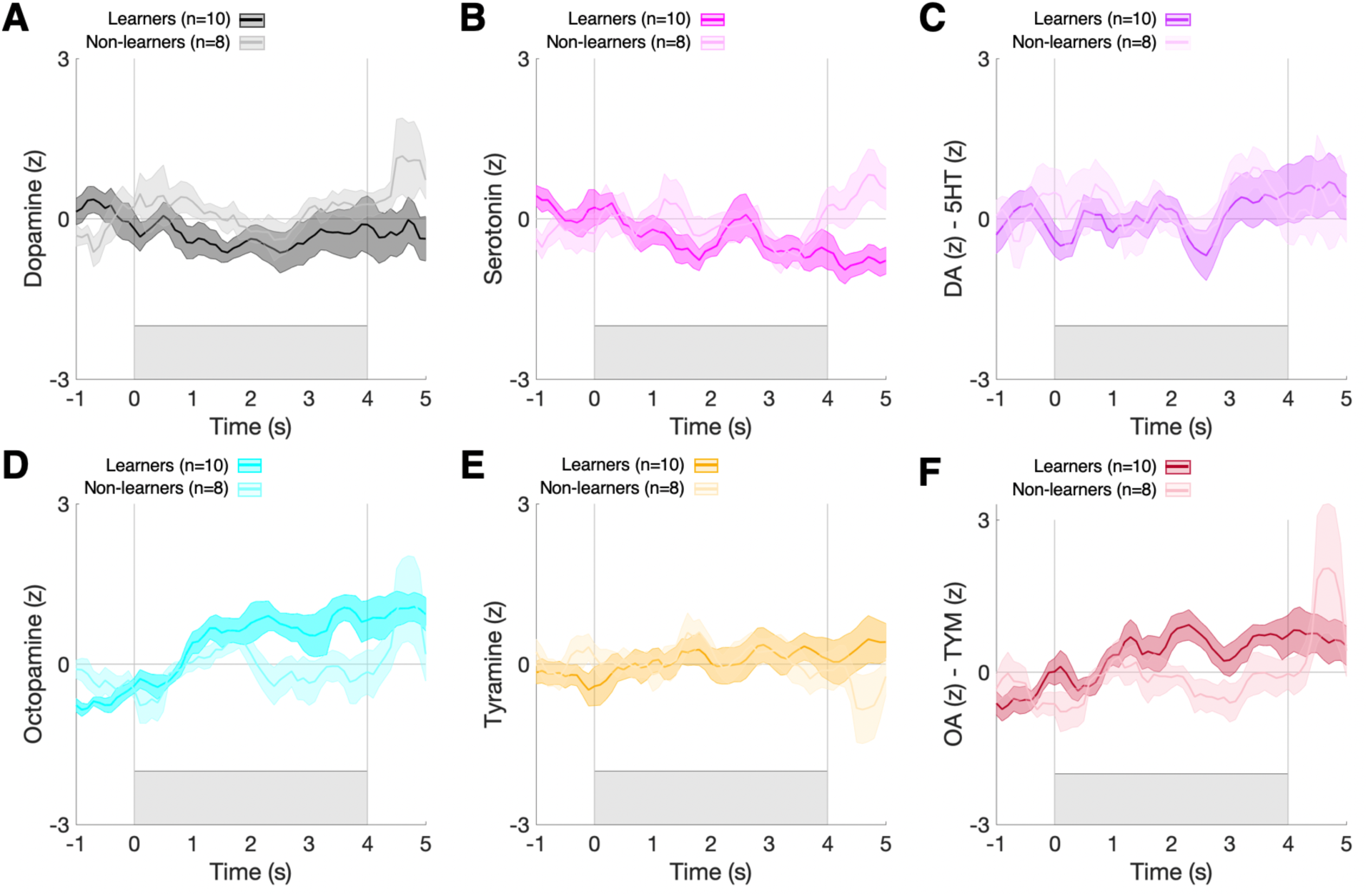
Learner and non-learner bee neurotransmitter response to hexanol exposure pre-conditioning. Average neurotransmitter and opponent pair time series in response to hexanol pre-conditioning for learner bees (n=10) and non-learner bees (n=8). Shaded regions indicate standard error of the mean (SEM), and the grey vertical lines and shaded rectangle indicate the hexanol temporal window.

**Fig. S3.**
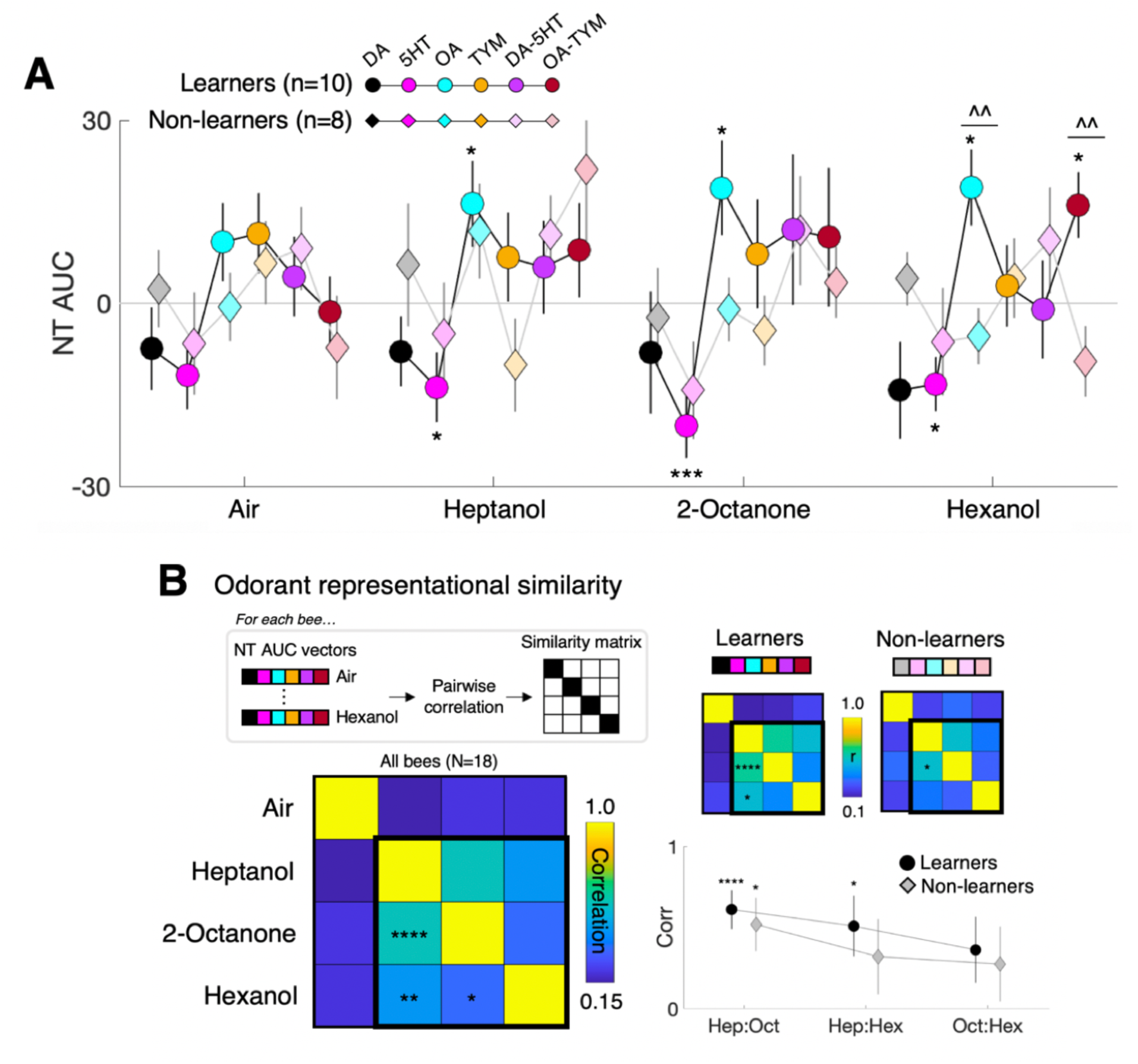
Learner and non-learner pre-conditioning odorant responses across neurotransmitters within antennal lobe. **(A)** Response levels (AUCs) for neurotransmitters and opponent pairs across the air and odor stimulation conditions. Circles and diamonds represent mean group-level AUC values, and the vertical lines depict SEM. *p<0.05,***p<0.005 one-sample t-test; ^^p<0.01 two-sample t-test. **(B)** Representational similarity analysis of neurotransmitter response patterns (vector of NT AUC values) to air, heptanol, 2-octanone, and hexanol exposure. Response levels (AUC) for DA, 5HT, OA, TYM, (DA-5HT), and (OA-TYM) time series in the 4-second odor presentation were correlated across all bees (left; N=18) as well as for learners and non-learners separately (right). Note that the matrices and color bars depict the random-effects representational similarity (i.e., computed for each bee individually). *p<0.05, **p<0.01, ****p<0.001 one-sample t-test.

**Fig. S4.**
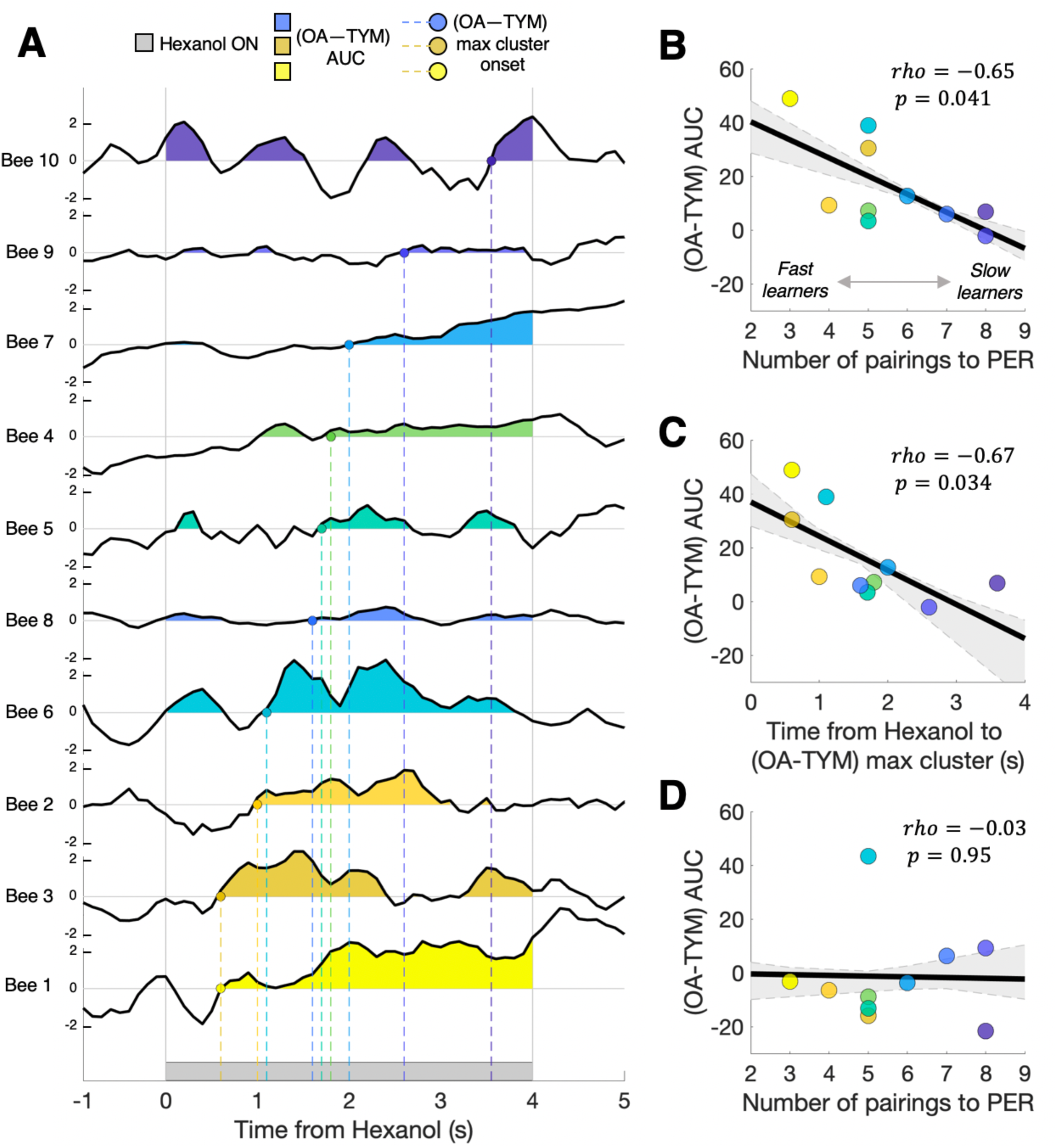
Learner bee (OA-TYM) response to first hexanol exposure pre-conditioning. **(A)** Individual (n=10) learner bees (OA-TYM) opponent response at the first hexanol exposure for each bee. Shaded regions indicate positive area under the curve, and the onset of the maximum positive cluster is indicated by the colored circles and dashed lines. **(B)** Correlation of individual learner bees’ (OA-TYM) AUC during hexanol exposure and their subsequent learning rate (trials to PER) during conditioning. **(C)** Correlation of individual learner bees’ (OA-TYM) max positive cluster onset time during hexanol exposure with their subsequent learning rate (trials to PER) during conditioning. **(D)** Correlation of learner bees’ (OA-TYM) AUC values during control air puff with their learning rate. In **(B-D)**, dashed lines and shaded region indicate the maximum and minimum of the leave-one-out bootstrapped correlation across learner bees.

**Fig. S5.**
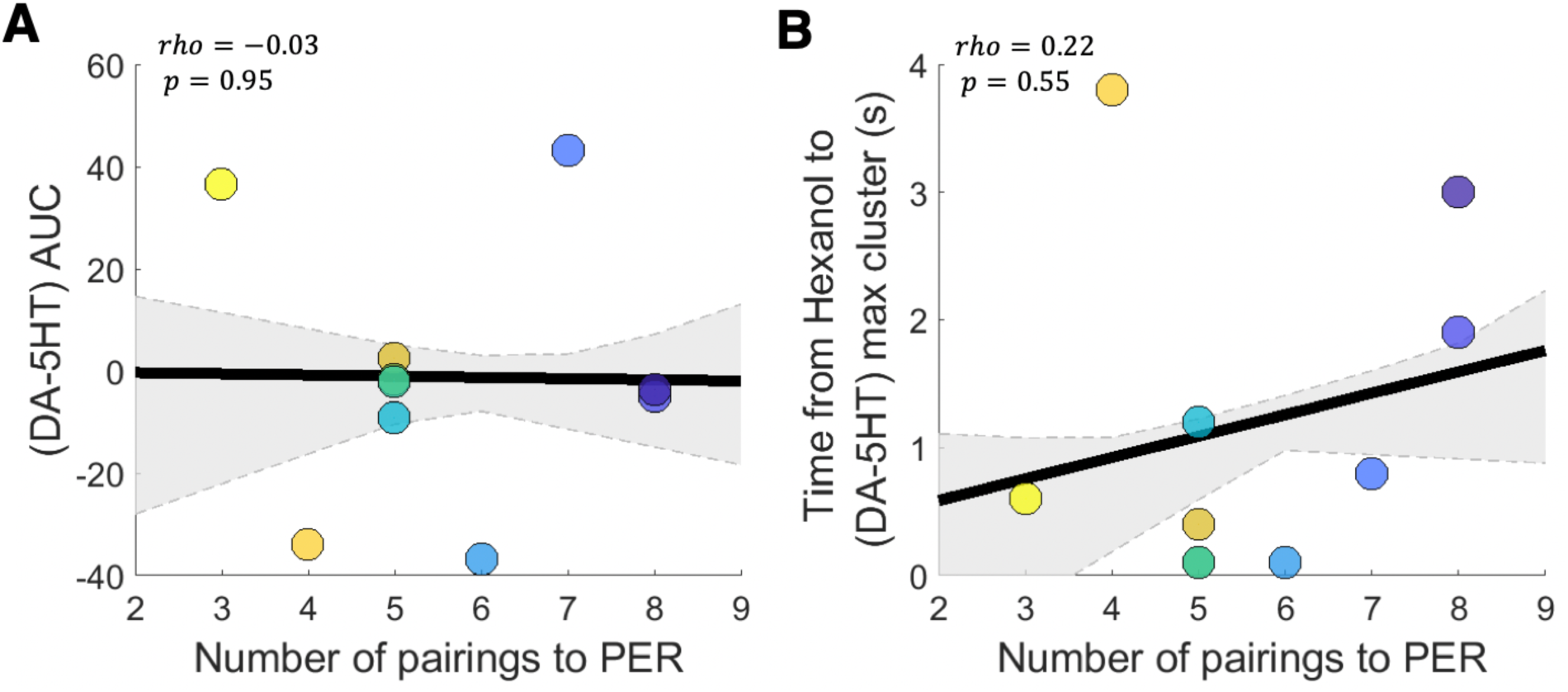
Learner bee (DA–5HT) response to first hexanol exposure pre-conditioning. **(A)** Correlation of individual learner bees’ (DA–5HT) AUC during hexanol exposure and their subsequent learning rate (trials to PER) during conditioning. **(B)** Correlation of individual learner bees’ (DA–TYM) max positive cluster onset time during hexanol exposure with their subsequent learning rate (trials to PER) during conditioning. In **(A,B)**, black line depicts the group-level correlation (rho), and the grey dashed lines and shaded region indicate the maximum and minimum of the leave-one-out bootstrapped correlation across learner bees.

**Fig. S6.**
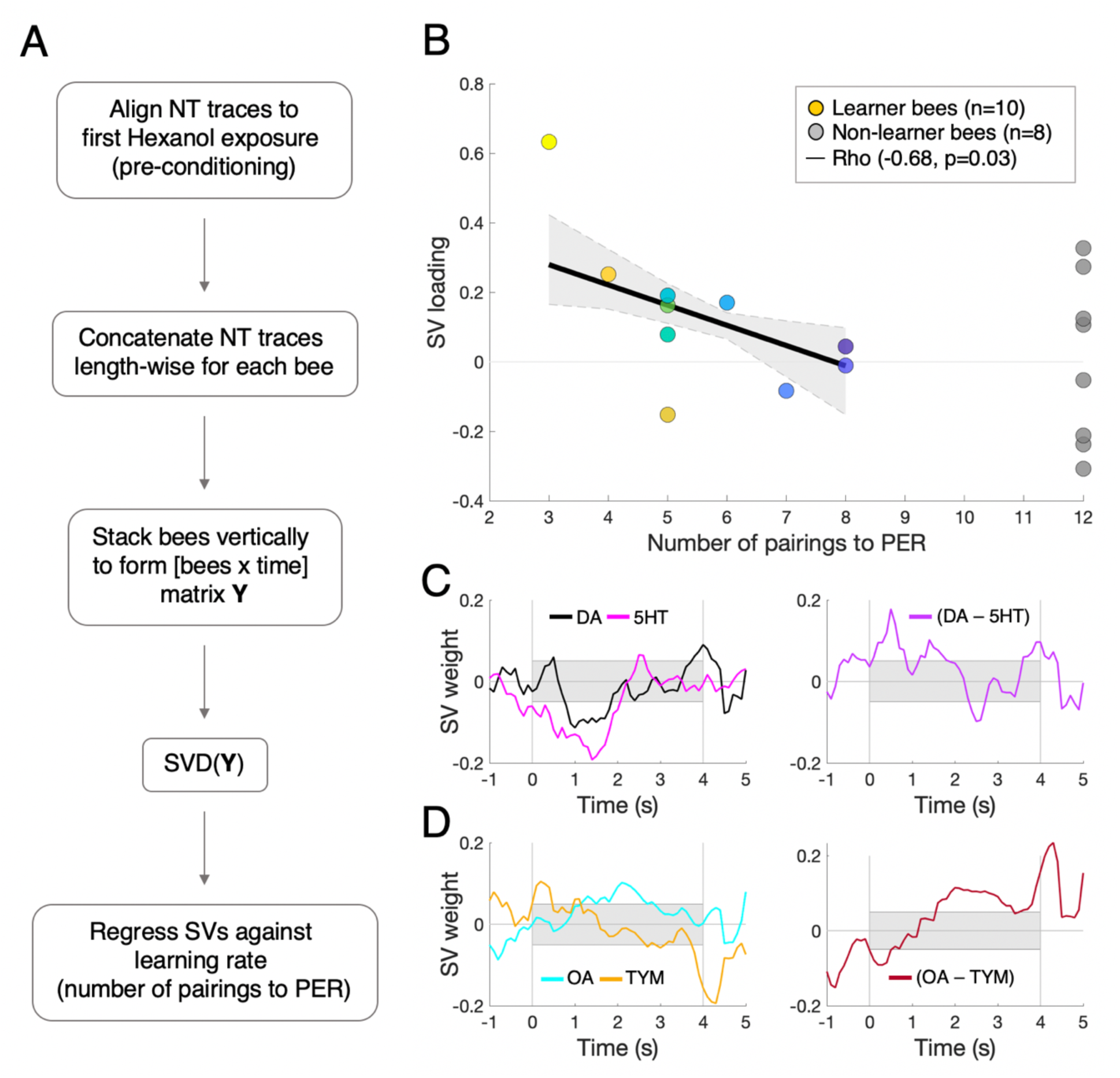
Singular value decomposition analysis. **(A)** Summary of SVD analysis steps (also see Methods). **(B)** Correlation of singular vector 4 loadings with learning rate (number of pairings to PER). **(C)** Latent pattern of dopamine (black) and serotonin (magenta) time series (left) and opponent response (right). Grey bar depicts presence of hexanol. **(D)** Same as **(C)** but for octopamine (cyan) and tyramine (orange).

**Fig. S7.**
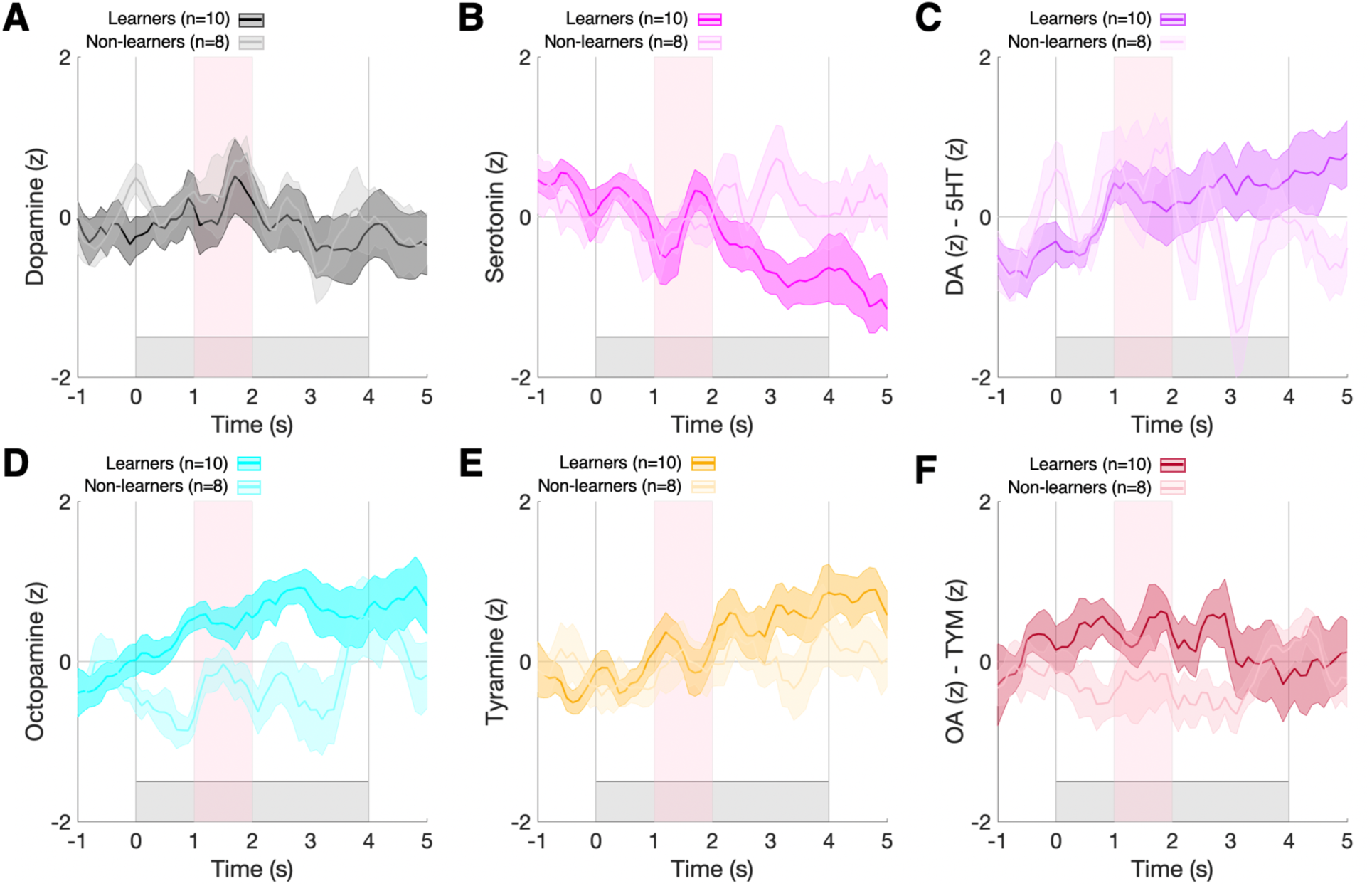
Learner and non-learner bee neurotransmitter response to first hexanol:sucrose pairing. Average neurotransmitter and opponent pair time series in response to the first hexanol:sucrose pairing during conditioning for learner bees (n=10) and non-learner bees (n=8). Shaded regions indicate standard error of the mean (SEM), and dashed lines indicate the air puff temporal window.

**Fig. S8.**
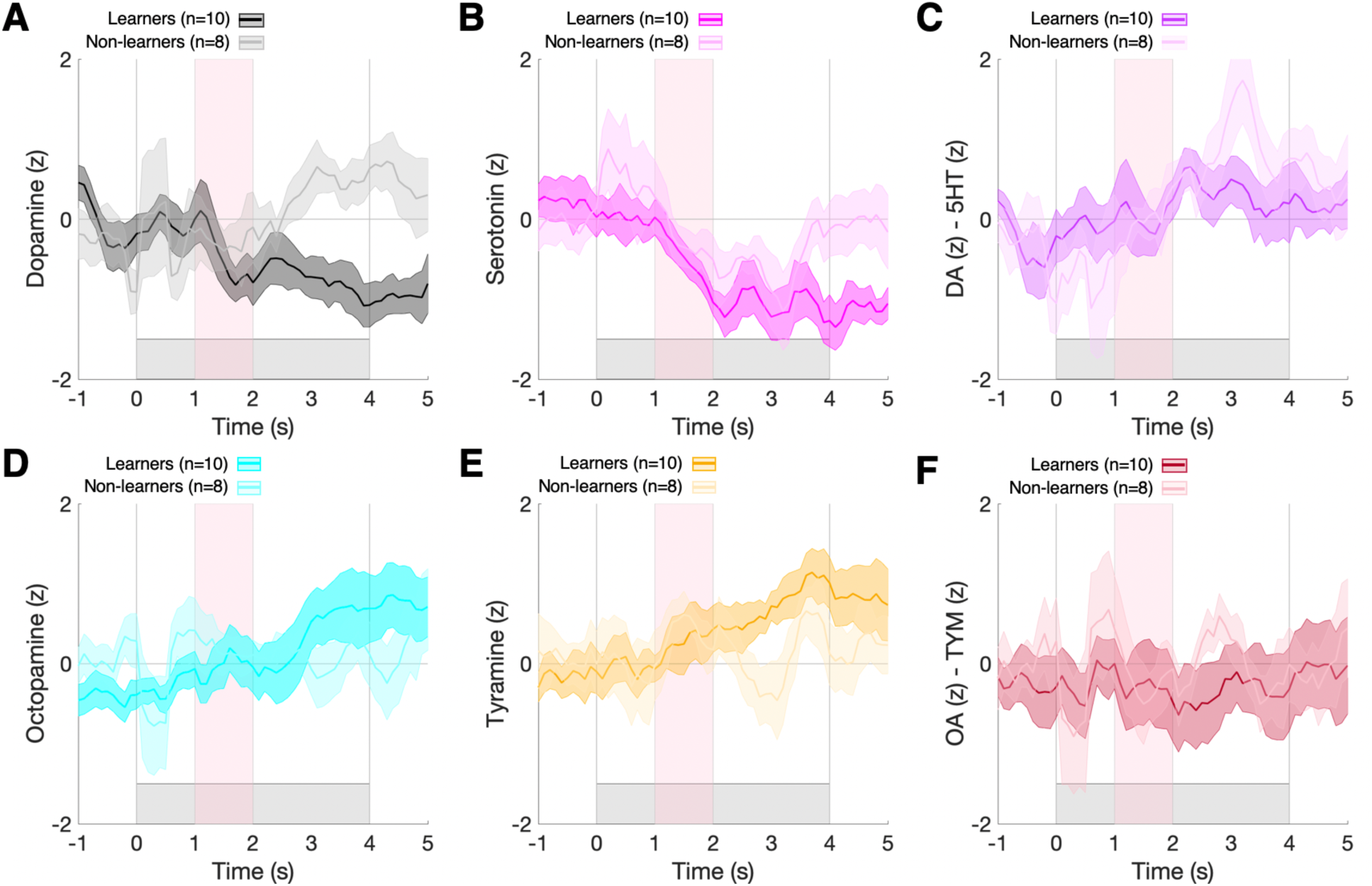
Learner and non-learner bee neurotransmitter response to last hexanol:sucrose pairing. Average neurotransmitter and opponent pair time series in response to last hexanol:sucrose pairing during conditioning for learner bees (n=10) and non-learner bees (n=8). Shaded regions indicate standard error of the mean (SEM), and dashed lines indicate the air puff temporal window.

**Fig. S9.**
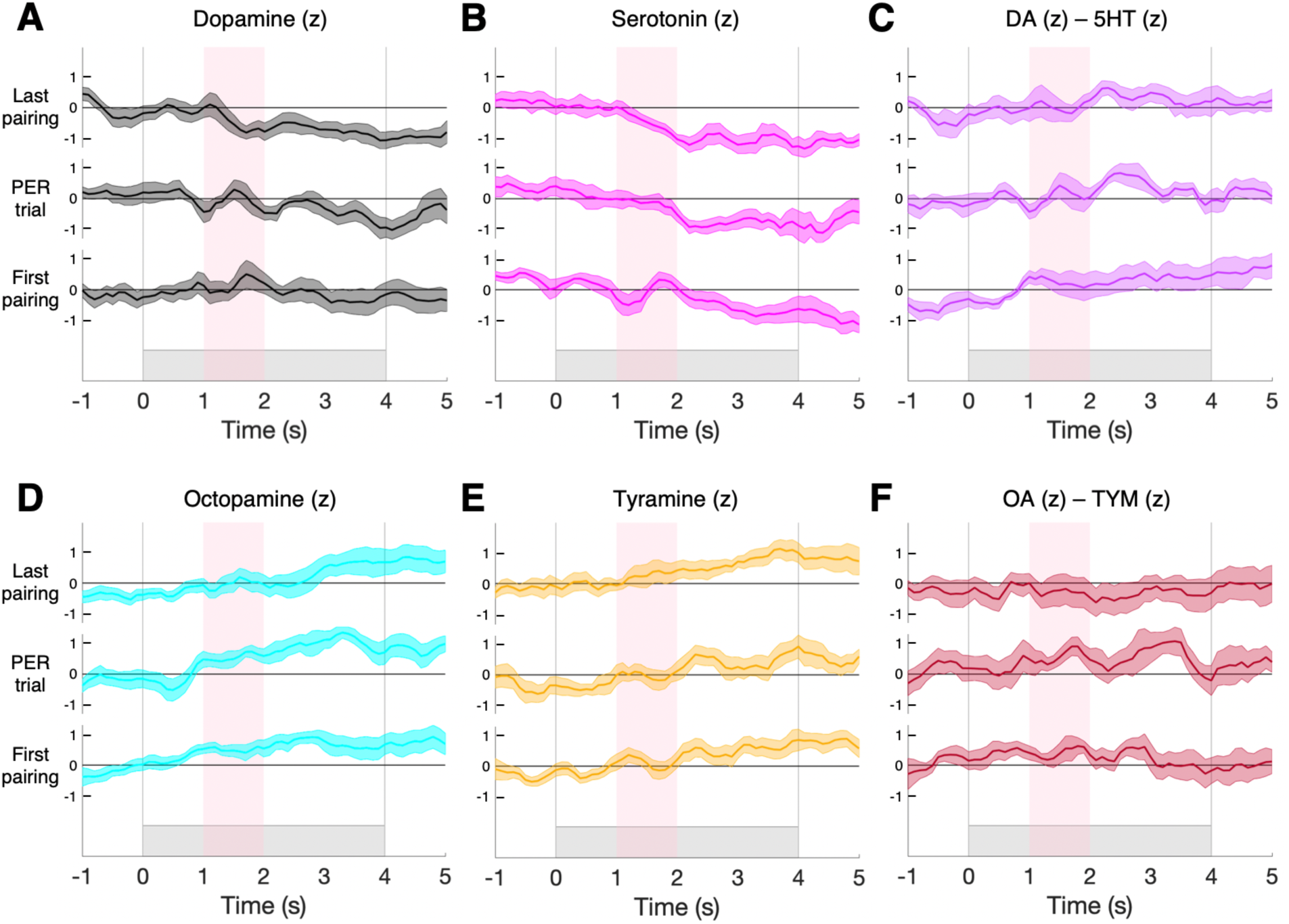
Learner bee neurotransmitter responses across conditioning. Average neurotransmitter and opponent pair time series in response to first hexanol:sucrose pairing, the PER trial, and the last hexanol:sucrose pairing during conditioning for learner bees (n=10). Shaded regions of time series indicate standard error of the mean (SEM); grey shaded region and vertical lines indicate the hexanol temporal window; the pink shaded region indicates sucrose delivery temporal window.

**Fig. S10.**
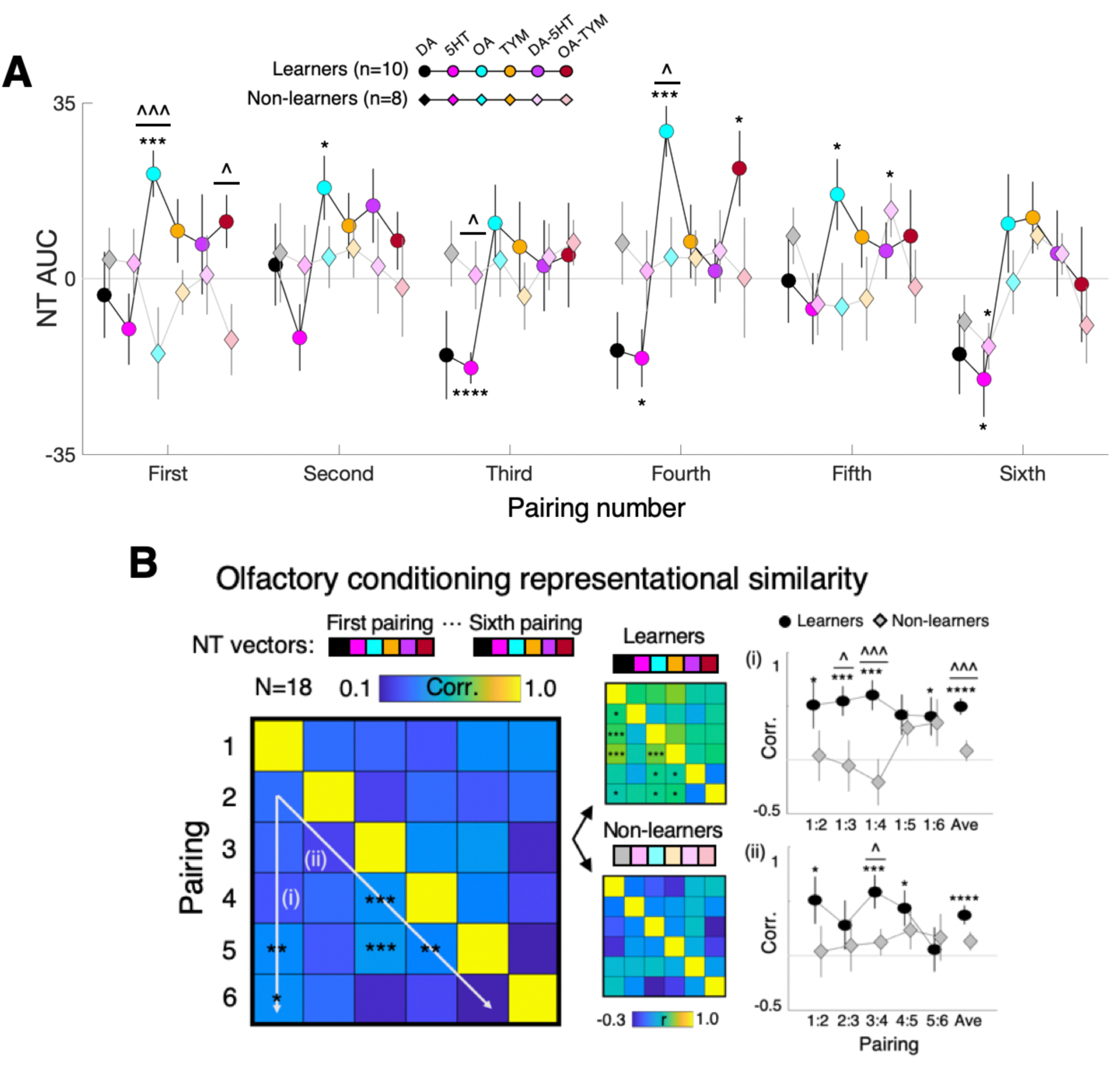
Learner and non-learner odorant conditioning responses across neurotransmitters within antennal lobe. **(A)** Response levels (AUCs) for neurotransmitters and opponent pairs for the first six CS:US pairings for learners and non-learners. Circles and diamonds represent mean group values; vertical lines depict SEM. *p<0.05, **p<0.01, ***p<0.005. ****p<0.001 one-sample t-test; ^p<0.05, ^^^p<0.005 two-sample t-test. **(B)** Representational similarity analysis of neurotransmitter response patterns to the first six CS:US pairings. Response levels (AUC) for DA, 5HT, OA, TYM, (DA-5HT), and (OA-TYM) time series in the 4-second odor presentation were correlated across all bees (left; N=18) and (right) for learners and non-learners separately. Note that the matrices and color bars depict the mixed-effects representational similarity (i.e., computed mean NT vectors for each bee individually). *p<0.05, **p<0.01, ***p<0.005, ****p<0.001 one-sample t-test. ^p<0.05, ^^^p<0.005 two-sample t-test.

**Fig. S11.**
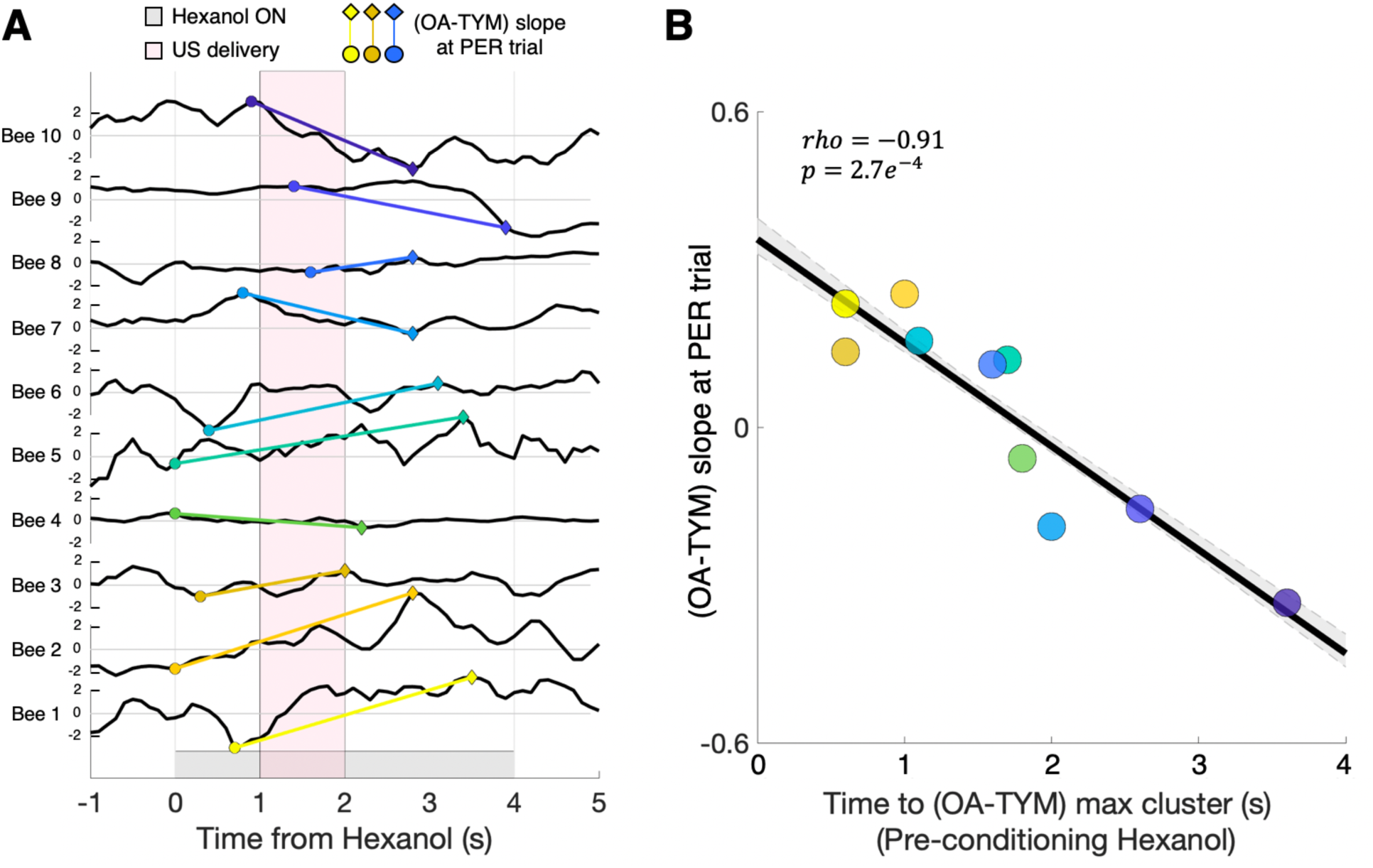
Learner bee (OA-TYM) rate of change at PER trial during odorant conditioning. **(A)** Individual (n=10) learner bees (OA-TYM) opponent response at the PER trial for each bee. Slopes are computed based on the extreme (max or min) response values in the first (0-2s; circles) and second half (2-4s; diamonds) of hexanol exposure. **(B)** Correlation of individual learner bees’ (OA-TYM) slope at PER trial and their pre-conditioning hexanol exposure (OA-TYM) max cluster onset timing.

**Table S1.**
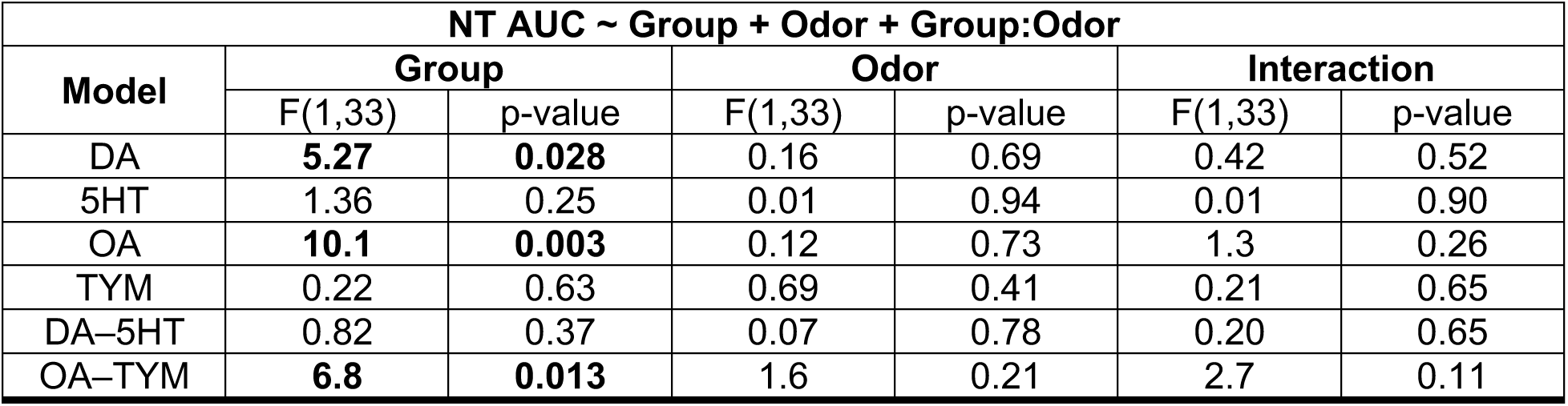
Linear model of pre-conditioning neurotransmitter responses to air and hexanol in all bees. The equation of the linear model is shown in the first row, with the ANOVA marginal statistical test results shown for each main effect of the model (group, odor) and their interaction between group and odor. For each ANOVA, the degrees of freedom are shown for the F-statistic.

**Table S2.**
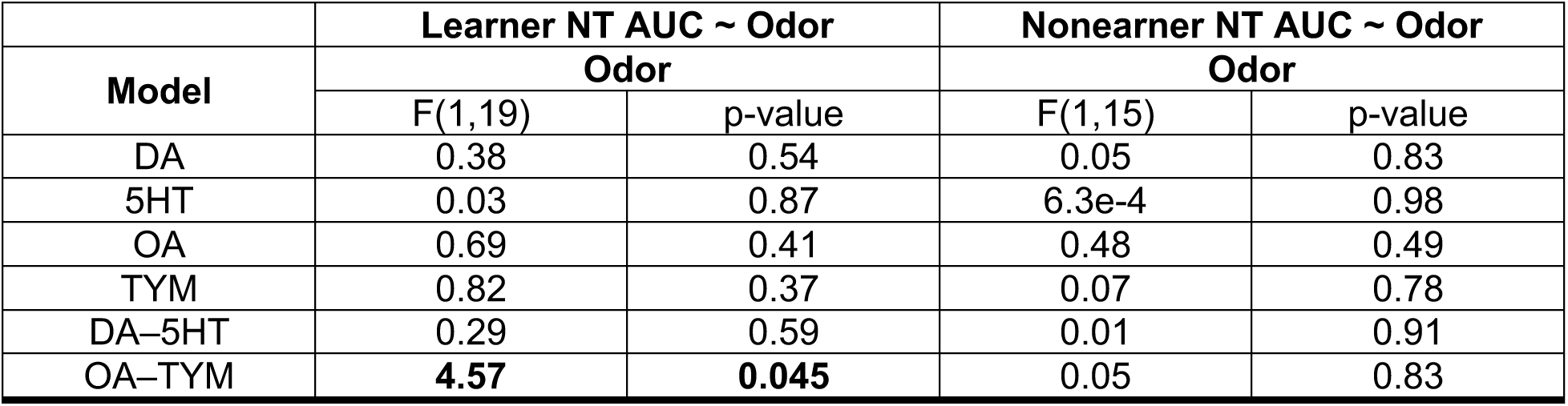
Linear model of pre-conditioning neurotransmitter responses to air and hexanol in learners and nonlearners. The equation of the linear model is shown in the first row, with the ANOVA marginal statistical test results shown for each main effect of the model (group, odor) and their interaction. For each ANOVA, the degrees of freedom are shown for the F-statistic.

**Table S3.** Linear model of neurotransmitter responses to first and last pairing during conditioning in all bees. The equation of the linear model is shown in the first row, with the ANOVA marginal statistical test results shown for each main effect of the model (group, pairing) and their interaction. For each ANOVA, the degrees of freedom are shown for the F-statistic.

| NT AUC ~ Group + Pairing + Group:Pairing |  |  |  |  |  |  |
| --- | --- | --- | --- | --- | --- | --- |
| Model | Group |  | Pairing |  | Interaction |  |
|  | F(1,33) | p-value | F(1,33) | p-value | F(1,33) | p-value |
| DA | **4.26** | **0.047** | 1.94 | 0.17 | 0.81 | 0.37 |
| 5HT | **5.94** | **0.020** | 3.55 | 0.07 | 0.05 | 0.82 |
| OA | **8.06** | **0.008** | 0.002 | 0.97 | **6.32** | **0.017** |
| TYM | **4.58** | **0.039** | 1.4 | 0.25 | 0.02 | 0.89 |
| DA-5HT | 0.098 | 0.75 | 0.17 | 0.68 | 0.31 | 0.58 |
| OA-TYM | 0.59 | 0.45 | 0.85 | 0.36 | 4.03 | 0.053 |

